## Supplementary material for "Recycling of cell surface membrane proteins from yeast endosomes is regulated by ubiquitinated Ist1": Combined supp material

### SUPPLEMENTAL FIGURES AND LEGENDS

- **Figure S1:** Differential trafficking itineraries of SNAREs and nutrient transporters
- **Figure S2:** The role of Ist1 mutants and potentially Npl4 in recycling
- **Figure S3:** Mass spec and pull-down controls to show Ist1 is ubiquitinated
- **Figure S4:** Assessments of Ist1 stability
- **Figure S5:** Ist1 localizations *in vivo*

### SUPPLEMENTAL MOVIES

- **Supplemental Movie S1**
  Mup1-GFP + Vps4-mCherry methionine pulse (short imaging intervals)
- **Supplemental Movie S2**
  Mup1-GFP + Sec7-mCherry methionine pulse (short imaging intervals)
- **Supplemental Movie S3**
  Mup1-GFP + Vps4-mCherry methionine pulse-chase (long imaging intervals)
- **Supplemental Movie S4**
  Mup1-GFP + Sec7-mCherry methionine pulse-chase (long imaging intervals)
- **Supplemental Movie S5**
  Mup1-mEos photoconversion methionine pulse-chase (long imaging intervals)
- **Supplemental Movie S6**
  Ist1-mCherry + Vps4-GFP time lapse imaging
- **Supplemental Movie S7**
  Ist1-mCherry + Mup1-GFP time lapse imaging

### SUPPLEMENTAL TABLES

- Supplemental Table T1: **Yeast Strains used in this study**
- Supplemental Table T2: **Plasmids used in this study**
- Supplemental Table T3: **Primary antibodies**
- Supplemental Table T4: **Statistical analyses**
- Supplemental Table T5: **Peptides identified by mass spectrometry**

### SUPPLEMENTAL REFERENCES

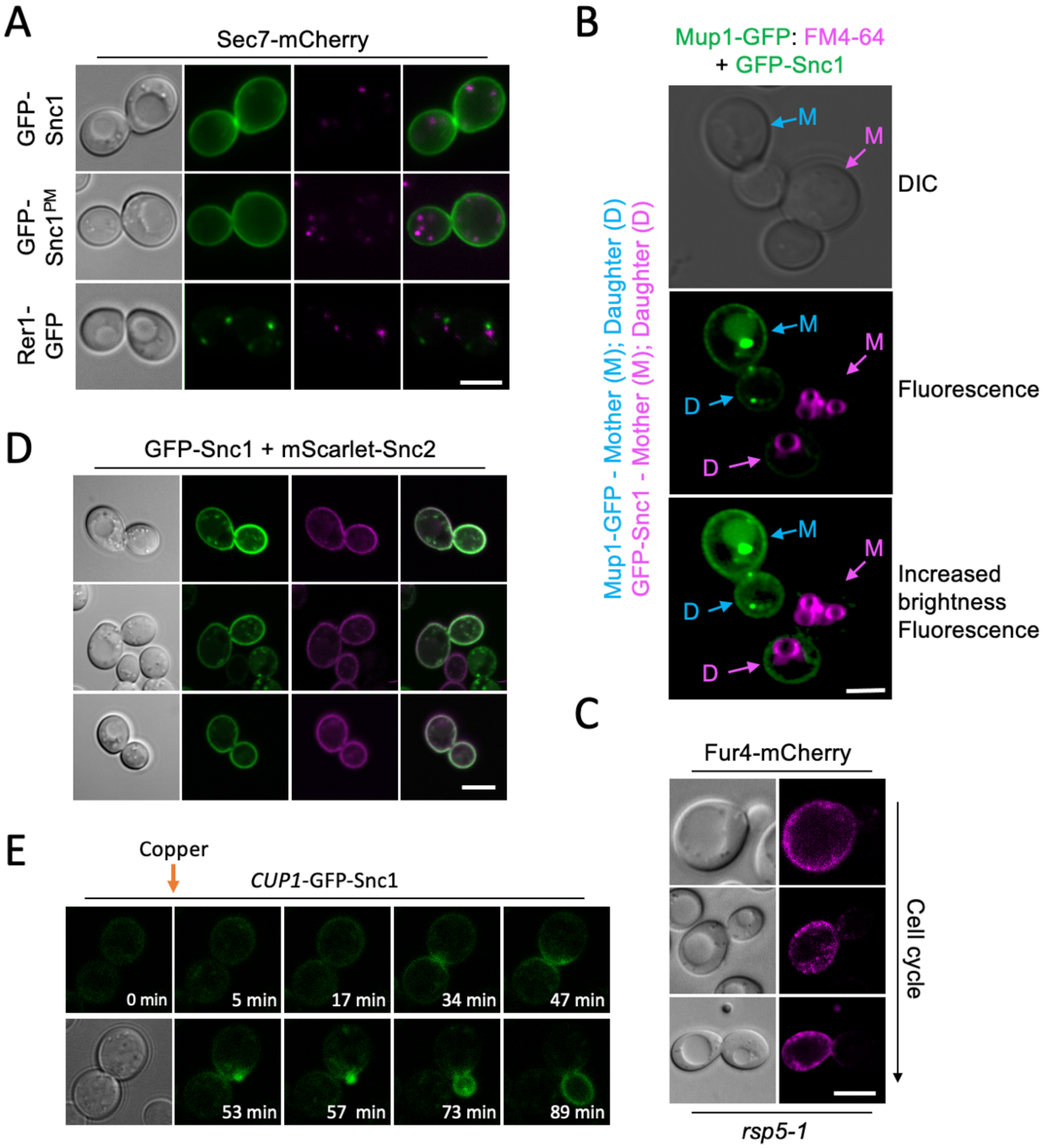


**Figure S1: Differential trafficking itineraries of SNAREs and nutrient transporters**
Airyscan confocal microscopy of wild-type cells expressing an endogenously expressed Sec7-Cherry and indicated GFP-tagged proteins plasmids show expected localisations. **B)** Wild-type cells expressing GFP-Snc1 were mixed with Mup1-GFP expressing cells previously pulse-chased with FM4-64 prior to imaging. **C)** *rsp5-1* cells expressing Fur4-mCherry were grown to mid-log phase in SC media and processed for confocal microscopy imaging. **D)** Airyscan microscopy shows colocalization and correct localisation of tagged versions of Snc1 and Snc2 expressed from the *CUP1* promoter. **E)** Expression of GFP-Snc1 following microfluidic addition of 20 μM copper chloride to cells during time-lapse microscopy. Scale bar, 5 μM.

A

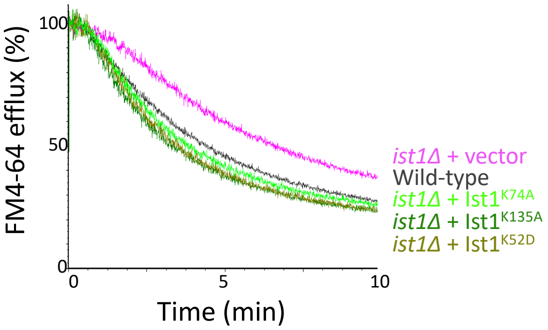


B

| Selected molecular function slim terms | | |
|---|---|---|
| **GO term** | **Description** | **Genes** |
| GO:0016787 | Hydrolase activity | *CDA2, HDA1, HOS2, NEM1, PTC1, RAS2,RPD3, SIT4, YNL010W, YVH1* |
| GO:0003924 | GTPase activity | *GPA1, GTR1, GTR2, RAS2, YPT11, YPT31, YPT32, YPT6* |
| GO:0019899 | Enzyme binding | *DEP1, **NPL4**, STE5, VAM6* |
| GO:0016887 | ATPase | *CDC50, DRS2, LEM3, VMA16* |
| GO:0032182 | Ubiquitin-like binding | ***NPL4*** |

### Figure S2: The role of Ist1 mutants and potentially Npl4 in recycling

**A)** Either wild-type cells or *ist1*Δ mutants transformed with vector (pink) or indicated Ist1 point mutations (green) were grown to mid-log phase prior to loading with dye for 8 minutes at room temperature, washing in cold media and efflux measured by flow cytometry. **B)** Table showing selected slim term annotations for recycling machinery associated with relevant enzyme activity that exhibit enrichment compared with genome-wide distribution.

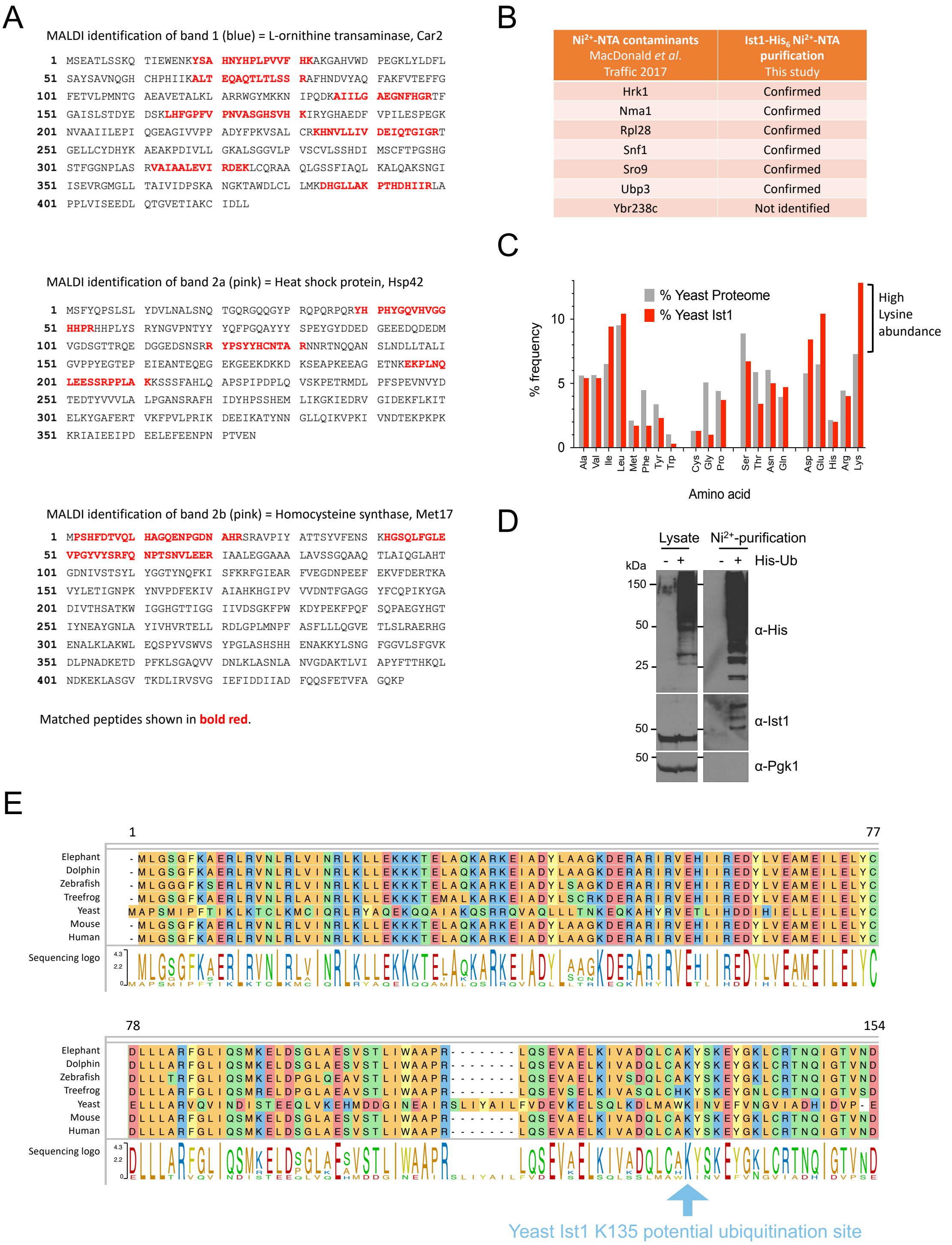


### Figure S3: Mass spec and pull-down controls to show Ist1 is ubiquitinated

**A)** Amino acid sequence of MALDI identified proteins Car2 (top), Hsp4 (middle), Met17 (bottom) with matched peptide sequences annotated in bold red . **B)** Table showing previously identified contaminants from yeast lacking $His_6$ tagged proteins purified on a $Ni^{2+}$-NTA column. Table also includes which contaminants were identified by mass spectrometry from purification of Ist1-HA- $His_6$. **C)** Histogram depicting average amino acid percentage distribution in yeast proteome (grey) and Ist1 (red). **D)** Wild t-type and $His_6$-ubiquitin expressing cells were grown to log-phase before ubiquitinated proteins were isolated from a denatured lysate on $Ni^{2+}$-NTA beads. Original lysates left and purified samples (right) were analysed by SDS-PAGE followed by immunoblot with the indicated antibodies. **E)** Ist1 amino acid sequence (from residue 1-154 from *saccharomyces cerevisiae* yeast) alignment across species indicated with the highly conserved lysine residue at position 135 (blue arrow.

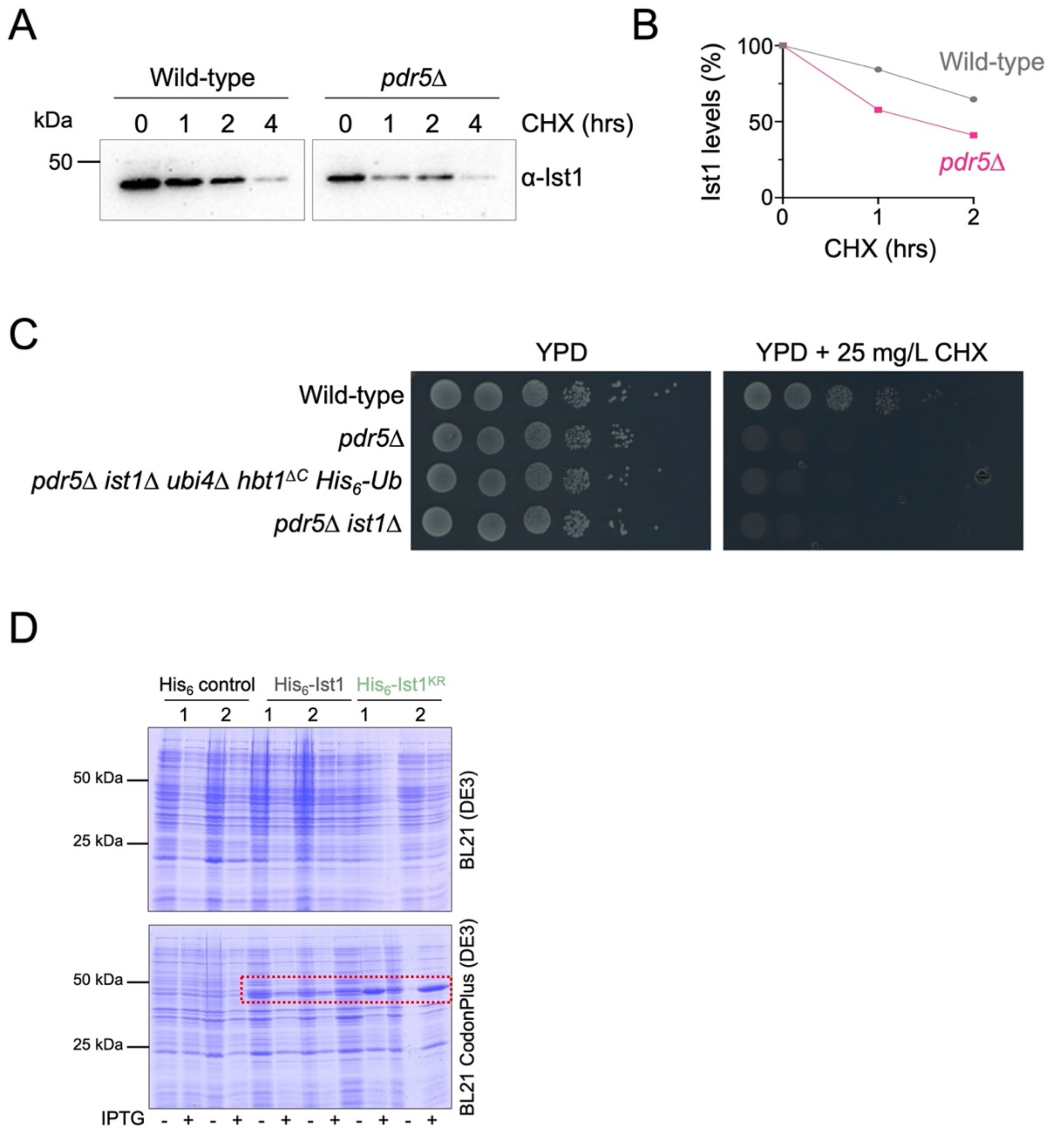


### Figure S4: Assessments of Ist1 stability

**A)** Cycloheximide chase experiments using wild-type and *pdr5Δ* cells grown to mid-log phase before exposure to 25 mg/L cycloheximide for denoted time before harvesting and immunoblot with α-Ist1 antibodies. **B)** Graph of Ist1 stability following cycloheximide chase at indicated times in wild-type (grey) and *pdr5Δ* (pink) cells. **C)** Growth assays in wild-type, *pdr5Δ*, *pdr5Δ ist1Δ hbt1*$^{\Delta C}$ *His*$_6$*-Ub*, and *pdr5Δ ist1Δ* yeast strains grown to exponentially dividing phase then spotted on YPD (left) and YPD containing 25 mg/L cycloheximide (right). **D)** SDS-PAGE Coomassie stained gels of lysates from BL21(DE3) (upper) or BL21 CodonPlus(DE3) (lower) *e.coli* strains expressing His$_6$ control, His$_6$-Ist1, or His$_6$-Ist1$^{KR}$ under *T7* promoter control. For each plasmid two clones are shown from lysates grown to $OD_{600}$=0.6 which were grown 16 hours at 15°C -/+ 0.5mM IPTG.

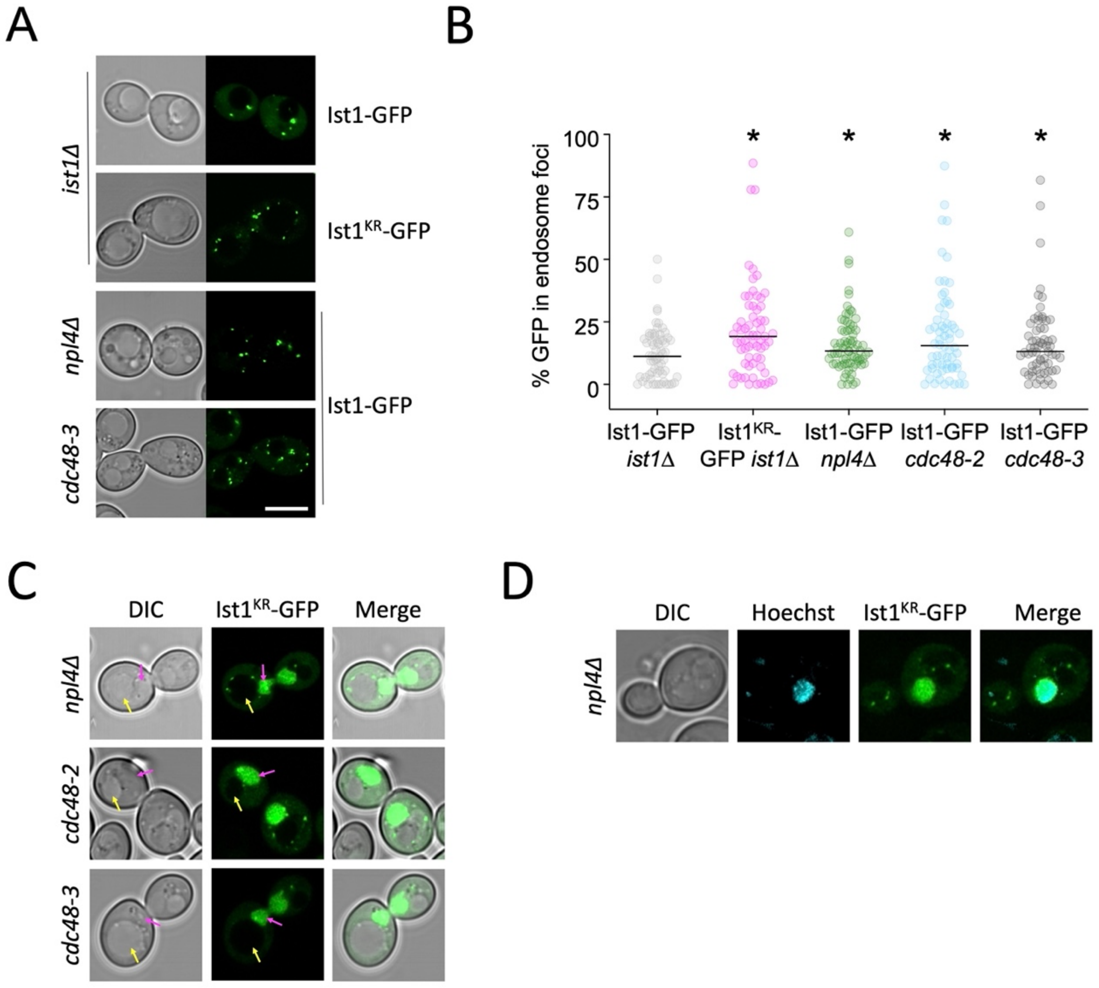


### Figure S5: Ist1 localizations *in vivo*

**A)** Indicated strains expressing Ist1$^{WT}$-GFP and Ist1$^{KR}$-GFP were imaged by Airyscan microscopy. **B)** The percentage GFP signal in endosomal foci quantified as a percentage of total cellular fluorescence. **C)** Ist1$^{KR}$-GFP mislocalizes to the nucleus (pink arrow or Hoechst stain) in indicated mutants. **D)** Airyscan microscopy used to localise indicated fluorescent proteins in wild-type cells. Scale bar, 5 μm.
